# iPromoter-BnCNN: a novel branched CNN based predictor for identifying and classifying sigma promoters

**DOI:** 10.1101/2019.12.27.884965

**Authors:** Ruhul Amin, Chowdhury Rafeed Rahman, Habibur Rahman Sifat, Nazmul Khan Liton, Moshiur Rahman, Sajid Ahmed, Swakkhar Shatabda

**Affiliations:** Department of Computer Science and Engineering, United International University, Dhaka, 1207, Bangladesh

## Abstract

**Motivation:** Promoter is a short region of DNA which is responsible for initiating transcription of specific genes. Development of computational tools for automatic identification of promoters is in high demand. According to the difference of functions, promoters can be of different types. Promoters may have both intra and inter class variation and similarity in terms of consensus sequences. Accurate classification of various types of sigma promoters still remains a challenge.

**Results:** We present **iPromoter-BnCNN** for identification and accurate classification of six types of promoters - *σ*^24^, *σ*^28^, *σ*^32^, *σ*^38^, *σ*^54^, *σ*^70^. It is a CNN based classifier which combines local features related to monomer nucleotide sequence, trimer nucleotide sequence, dimer structural properties and trimer structural properties through the use of parallel branching. We conducted experiments on a benchmark dataset and compared with six state-of-the-art tools to show our supremacy on 5-fold cross-validation. Moreover, we tested our classifier on an independent test dataset.

**Availability:** Our proposed tool iPromoter-BnCNN web server is freely available at http://103.109.52.8/iPromoter-BnCNN. The runnable source code can be found here.

**Supplementary information:** Supplementary data (benchmark dataset, independent test dataset, model files, structural property information, attention mechanism details and web server usage) are available at *Bioinformatics*. online.

## 1 Introduction

Promoters are small regions near gene containing 100 to 1000 base-pairs. For transcription occurrence, RNA polymerase must bind near the promoter. Bacteria with prokaryotic cell type has promoters consisting of a purine at the transcription start site (TSS). It contains specific hexamers centered at -10 and -35 (Busby and Ebright (1994), Feng *et al.* (2017)). There are several sigma factors in the RNA polymerase of Escherichia coli bacteria, which are dependent on environment and gene. As a result, sigma factors are used as distinguishing elements of promoter sequences found in DNA. Each of the six different types of sigma factors such as *σ*^24^, *σ*^28^, *σ*^32^, *σ*^38^, *σ*^54^, *σ*^70^ has different functions. For example, *σ*^70^ factor is responsible for transcription of most of the genes under normal condition (Gruber and Gross (2003)). On the other hand, *σ*^24^ factor is responsible for heat shock response (Raina *et al.* (1995)). Similarly, *σ*^28^, *σ*^32^, *σ*^38^ and *σ*^54^ are responsible for flagellar genes, heat shock response, stress response during the transition from exponential growth phase to the stationary phase of E. coli (Jishage and Ishihama (1995)) and nitrogen metabolism, respectively (Janga and Collado-Vides (2007)).

Molecular techniques for promoter identification or classification is costly in terms of time and money, which is why computational methods are more popular (Towsey *et al.* (2008)). Promoters normally differ from the consensus at one or more positions. So, it is challenging to precisely predict promoters through traditional methodology.

Recently, a few computational methods have been proposed to classify DNA sequences as promoters or non-promoters, some aiming at identifying a certain class of sigma promoters. For instance, Coelho *et al.* (2018) provided BacSVM+, a software package using LibSVM library for promoter prediction in Bacillus subtilis. Work of Scheila de Avila e Silva* (2014) integrated DNA duplex stability as feature of neural network to identify *σ*^28^ and *σ*^54^ class of promoter in E. coli bacteria. Lin *et al.* (2014) developed iPro54-PseKNC which performs the same task using SVM classifier based on pseudo k-tuple nucleotide composition (PseKNC). Li *et al.* (2015) applied a deep feature selection (DFS) model on enhancer-promoter classification. Lin *et al.* (2017) used pseudo nucleotide composition for feature extraction in order to identify *σ*^70^ promoters in prokaryotes using SVM. He *et al.* (2018) used PSTNPSS(Position-specific trinucleotide propensity based on single-stranded characteristic) and PseEIIP(Electron-ion potential values for trinucleotides) features while Rahman *et al.* (2019a) used multiple windowing and minimal features for the same task. Rahman *et al.* (2019b) developed iPromoter-FSEn for performing the same task using feature subspace based ensemble classifier achieving an impressive accuracy of 86.32%. Umarov and Solovyev (2017) trained CNN based architecture on the same promoter type in E. coli. Liu *et al.* (2017) developed iPromoter-2L which can identify promoter and can classify them into six types. They used random forest PseKNC. Zhang *et al.* (2019) proposed MULTiPly for the same task using sequence-specific local information (k-tuple nucleotide composition, dinucleotide based autocovariance), bi-profile Bayes and KNN feature encoding schemes which incorporate position-specific residue distribution from the full set of training sequences and the distribution of promoters and non-promoters in the vicinity of the sequences, respectively. They applied F-score feature selection method to identify feature from each category giving the best prediction results.

Shahmuradov *et al.* (2017) worked on predicting transcription start sites (TSSs) in five types of E. coli sigma promoters such as - *σ*^24^, *σ*^28^, *σ*^32^, *σ*^38^ and *σ*^70^ though they did not work on E. coli sigma promoter classification. Only Liu *et al.* (2017) and Zhang *et al.* (2019) proposed computational methods (iPromoter-2L and MULTiPly) for classifying sigma promoters into six classes in E. coli bacteria. The sensitivity and specificity of promoter classification showed opposing behavior for iPromoter-2L. For example, for *σ*^28^, *σ*^32^, *σ*^38^ and *σ*^54^, iPromoter-2L showed specificity of higher than 99%, but the sensitivity was lower than 54%. The promoter classification performances of the binary sub-classifiers used in MULTiPly were impressive. For example, the first sub-classifier showed 85.24% accuracy in *σ*^70^ promoter type identification. The sensitivity and specificity were 87.27% and 86.57%, respectively. We follow the stage by stage binary classification method used in MULTiPly in this work. The main limitation of MULTiPly was the selection of the basic features to work with. Different combination of different heterogeneous features led to different prediction results.

We propose iPromoter-BnCNN, a one dimensional CNN based classifier which can identify sigma promoter and can classify sigma promoter into the six specified classes in E. coli bacteria. Four parallel branches of one dimensional convolution filters learn and and extract important local features related to monomer, trimer nucleotide sequence and dimer, trimer structural properties simultaneously. Dense layers at the end of our designed model combine these extracted features and perform the classification task. We compare our method with state-of-the-art tools for E. coli sigma promoter identification and classification and show the effectiveness of our method.

## 2 Materials and Methods

We followed Chou’s five-step rules (Chou (2011)) for more effective presentation of our research work. A series of recent publications (Xu *et al.* (2013), Xu *et al.* (2014), Liu *et al.* (2015), Jia *et al.* (2016a), Chen *et al.* (2016), Feng *et al.* (2017), Liu *et al.* (2017), Zhang *et al.* (2019)) comply with this standard. Briefly, the five steps are: (i) valid benchmark dataset selection (ii) biological sequence sample formulation with mathematical expression (iii) powerful algorithm introduction for prediction purpose (iv) predictor performance evaluation using cross validation (v) public access establishment to the constructed predictor. Our system overview has been provided in Figure 1 in light of the five steps described.

**Fig. 1.**
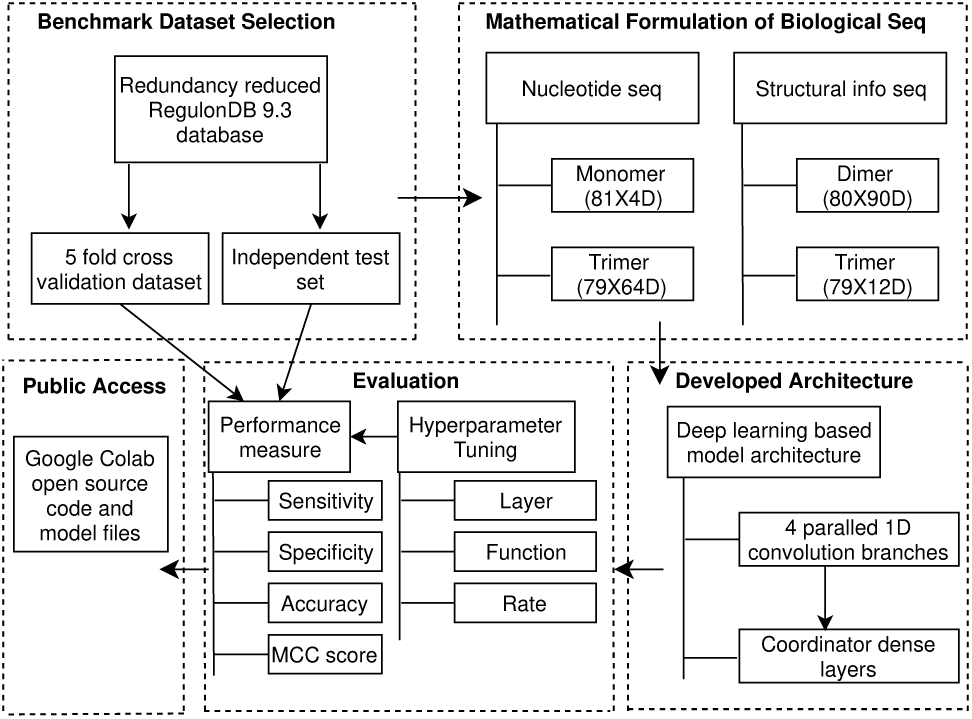
System overview of iPromoter-BnCNN

### 2.1 Benchmark Dataset

One benchmark dataset is good enough to prove the effectiveness of a certain method when K-fold cross-validation is used, because such evaluation takes into account the results obtained from K number of disjoint training and validation sets (Chou and Shen (2007)). We have used the same E. coli bacteria promoter dataset as of Liu *et al.* (2017) and Zhang *et al.* (2019) for comparison purpose in terms of sigma promoter identification and classification into sub types. All promoter samples of the used dataset are experimentally verified (each has 81 bp). They have been collected from the RegulonDB database (Version 9.3) (Gama-Castro *et al.* (2016)). Lin *et al.* (2014, 2017) randomly extracted non-promoter sequences from middle regions of long coding sequences and convergent intergenic regions in E.coli K-12 genome, which are also 81 bp long. We include these sequences in the non-promoter class. We have ensured redundancy reduction (no two samples of same class with pairwise sequence identity *≥* 0.8) using CD-HIT software (Li and Godzik (2006)) on our dataset following Liu *et al.* (2017) and Zhang *et al.* (2019). We use some recently included promoter samples (experimentally verified) from RegulonDB version 10.7 (Santos-Zavaleta *et al.* (2019)) as our independent test dataset. Our benchmark dataset and test dataset are disjoint. Dataset information have been provided in Table 1.

**Table 1:** Class-wise sample numbers in datasets used

| Classes | Benchmark Dataset | Independent Test Dataset |
| --- | --- | --- |
| Promoter | 2860 | 256 |
| Non-Promoter | 2860 | 0 |
| $\sigma^{24}$-promoter | 484 | 30 |
| $\sigma^{28}$-promoter | 134 | 4 |
| $\sigma^{32}$-promoter | 291 | 13 |
| $\sigma^{38}$-promoter | 163 | 10 |
| $\sigma^{54}$-promoter | 94 | 0 |
| $\sigma^{70}$-promoter | 1694 | 199 |

### 2.2 Mathematical Formulation of DNA Sequence

Computational methods require effective numerical vector representation of nucleotide sequences for prediction or classification tasks (Chou (2015)). We consider vector representation of two categories for our work. We describe them in the following subsections.

#### 2.2.1 Original Nucleotide Sequence

A DNA sequence can be expressed as follows:

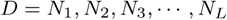

where, *L* is the length of the DNA sequence and *N*_*i*_ ∈ {*A, T, C, G*}. Although a DNA sequence comprising of nucleotides do not show any distinguishing property when looked at visually, deep learning based models are powerful enough to infer various distinguishing features from local patterns if we can represent such sequence with appropriate mathematical representation (Umarov and Solovyev (2017), Singh *et al.* (2016), Xu *et al.* (2016)). In a DNA sequence, there can be four types of monomers such as - A, T, C and G. So, our monomer representation of each DNA sample is a 81 × 4 size two dimensional matrix (each sequence is 81 nucleotide long in our dataset). Each nucleotide is represented by a one hot vector (1 in one position, all other positions 0) of size four.

We also construct an overlapping trimer representation of each DNA sequence. Each codon corresponding to a single amino acid is a combination of three nucleotides. Full set of codons form the genetic code. This is why trimers have special significance. In the *L* length DNA sequence mentioned in this subsection, there are total *L* − 2 overlapping trimers which are as follows:

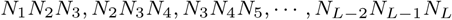

There can be 4^3^ = 64 kinds of possible trimers. We represent each trimer with a one hot vector of size 64. Thus, each DNA sample is represented by a 79 × 64 size two dimensional matrix.

#### 2.2.2 Structural Properties

Structural property refers to specific characteristics of DNA molecule such as stability, rigidity or curvature (Meysman *et al.* (2012)). Conformational properties are related to static DNA structure (geometrical property) while physicochemical properties are related to dynamic DNA structure (potential to change in conformation). These properties play an important role in promoter prediction and classification (Abeel *et al.* (2008), Bansal *et al.* (2014)). Chen *et al.* (2014b) constructed PseKNC-General tool which can convert DNA sequence dataset into pseudo nucleotide compositions providing many choices of physicochemical combinations. This tool provides 90 physicochemical properties (role, twist, tilt etc) for each of the 16 possible dimers and 12 physicochemical properties (trinucleotide GC content, consensus role, consensus rigid etc) for each of the 64 possible trimers. We implement physicochemical property wise normalization (subtract mean and divide by standard deviation) so that each property gets equal chance to act as distinguishing property. We provide the values of these physicochemical properties associated with each type of dimers and trimers as part of the supplementary information. In *L* length DNA sample, there are *L* − 1 overlapping dimers such as *N*_1_*N*_2_, *N*_2_*N*_3_, *N*_3_*N*_4_, *…, N*_*L*−1_*N*_*L*_. We replace each of these dimers with the 90 physicochemical properties and get a 80 × 90 size two dimensional matrix for each 81 length DNA sequence sample. Similarly, there are total *L* − 2 overlapping trimers such as *N*_1_*N*_2_*N*_3_, *N*_2_*N*_3_*N*_4_, *N*_3_*N*_4_*N*_5_, *…, N*_*L*−2_*N*_*L*−1_*N*_*L*_. We replace each of these trimers with the 12 physicochemical properties and get a 79 × 12 size two dimensional matrix for each 81 length DNA sequence sample.

### 2.3 Model Architecture

We use four kinds of feature representations for our model - monomer sequence matrix, trimer sequence matrix, dimer physicochemical property matrix and trimer physicochemical property matrix (two dimensional) of dimension (81, 4), (79, 64), (80, 90) and (79, 12), respectively as described in Subsection 2.2. We provide our model architecture in Figure 2. Each of these four unique representations of a sample sequence is passed through a separate one dimensional convolutional neural network (1D CNN) branch parallely as shown in the figure.

**Fig. 2.**
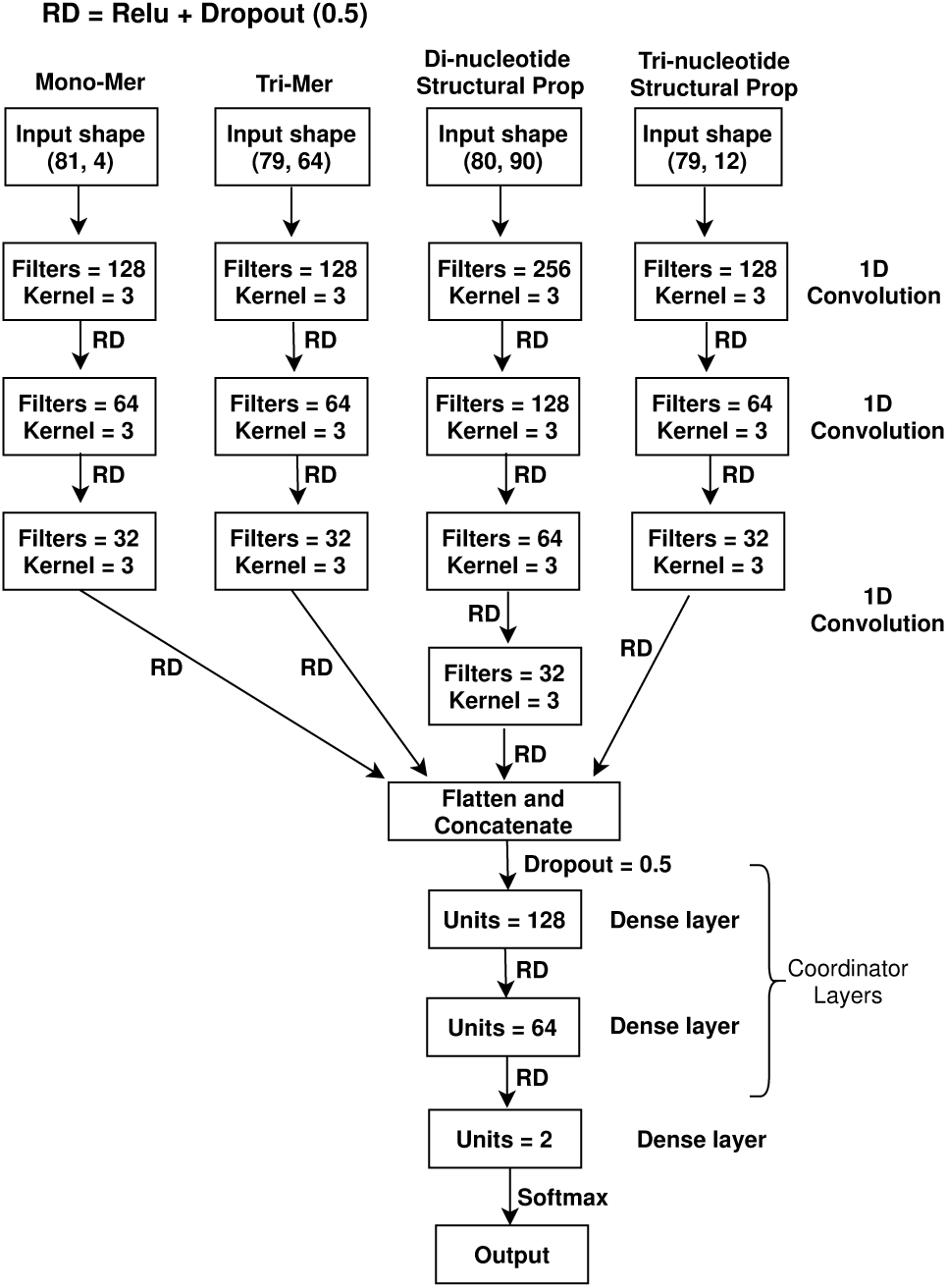
Model Architecture

1D CNN has shown its potential and significance in recent studies (Chen *et al.* (2017), Zhou *et al.* (2015), Oh *et al.* (2018)) related to local feature extraction and sequence data classification when the positions of the existing local features are not important. Each of the four branches of 1D CNN works as automatic distinguishing feature extractors for our classification task. The leftmost branch of Figure 2 gets us distinguishing sequence motifs from constituent monomers. The second branch learns locally important combination of codons. The third branch extracts distinctive patterns related to structural properties of the sequences based on numerical values that depict the contribution from each of the constituent dimers to these properties. The fourth branch performs the same task, but only this time from values representing constituent trimer contributions.

Each of the four different branches learns important distinguishing features from local sequence patterns. In order to perform a successful classification, we need to have a way to combine these independently learnt and extracted features. Each of these four branches return a matrix (two dimensional) of dimension *m*_*i*_ × *n*_*i*_, where the branch number is *i*. We flatten each of these matrices into a one dimensional vector and concatenate all four of them. The resultant one dimensional concatenated vector is of length 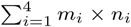. This now works as a feature vector. Instead of using this feature vector directly for classification, we pass this vector through densely connected neural network layers so that our model is able to learn successfully the importance of each feature for the classification task at hand. While the flattened output vectors from the four parallel branches can be regarded as independent feature groups, the coordinator layer (labeled in Figure 2) serves as an implicit feature selector and high-level attribute extractor that operates on an amalgamation of the feature groups.

Our representation scheme preserves the order of the nucleotide residues in which they appear in the DNA sequence. This can be exploited by the convolutional filters in different layers for extracting sequence-order information if deemed necessary for ensuring maximum separation between regions of interest. The initial filters that interact directly with the input layer can hypothetically encode information similar to *k*-tuple nucleotide composition, which is efficacious for representing DNA sequences in various property and function modeling tasks, including promoter identification (Zhang *et al.* (2019)). Additionally, the final layers of our model can encode long-range sequence order information due to the hierarchical output-input structure of the network, which allows these high-level filters to widen their receptive fields substantially. These filters can serve as a surrogate for features like pseudo *k*-tuple nucleotide composition (Chen *et al.* (2014b)) and gapped k-mers (Muhammod *et al.* (2019)) which capture useful long-range sequence order information from structural profile and primary sequence, respectively. Due to these hierarchical interactions between initial and high-level filters, our architecture can also extract information like GC content (indicates proportion of G and C bases out of four total bases in a genome region) in particular regions of the input sequences, which has been quite successful in promoter identification (Abeel *et al.* (2008)). Preservation of residue-order is further facilitated by the architectural choice of not adding any pooling layers which seemed to degrade performance in our initial experiments due to loss of spatial information. Furthermore, pooling layers may disrupt structural property extraction from di and tri nucleotide contribution scales since these properties are usually contingent on the position-specific neighborhood structure of the nucleotides in the original sequence (Meysman *et al.* (2012)).

Although opting for a feature extractor based on a neural network that does not incorporate any unidirectional or bidirectional recurrent layers may seem a little counter-intuitive in the context of biological sequence modeling, this specific modeling choice has ensured robustness of the extracted features by keeping the number of learnable parameters comparatively small through parameter sharing. Such simplified feature transformation functions are of paramount importance for curbing idiosyncratic pattern extraction given our relatively small-scale training data, which is often the case for new genome projects (Abeel *et al.* (2008)). Additionally, we have opted for comparatively shallow convolution branches for keeping the sequence to promoter-class mapping function as simple as possible without losing the required feature extraction capability of the model.

We use **relu** as activation function for each intermediate layer as it has become very popular for its simplicity and effectiveness (Li and Yuan (2017), Yarotsky (2017), Agarap (2018)). In the final layer, we use softmax function for binary classification purpose with two nodes constituting the last dense layer. Since our training data is relatively small, we have employed dropout regularization, which ignores some layer outputs randomly. Such treatment changes the connectivity of a layer with its previous layer on each epoch of training forcing the model architecture to look different every time (Srivastava *et al.* (2014), Srivastava (2013), Baldi and Sadowski (2013)). We use a high dropout rate of 0.5 (output from a particular layer node is ignored with 50% probability) after each of our layers (except for the output layer) to prevent overfitting.

### 2.4 Model Selection and Performance Evaluation

The goal of iPromoter-BnCNN is to identify a query DNA sequence as a promoter or non-promoter and in case of being promoter, to predict which of the six types of promoters the identified promoter belongs to. Although promoter classification is a multi-class classification problem, the dataset that we use has severe class imbalance problem. For example, there are 1694 samples in *σ*^70^-promoter, the largest promoter subset while only 94 samples belong to the smallest promoter subset *σ*^54^-promoter. Our deep learning based model showed poor training performance when we used the smart undersampling technique introduced for training bTSSfinder tool by Shahmuradov *et al.* (2017). The probable reason is that the minority class *σ*^54^ has only 94 samples and such low number of samples from each class is not enough to train deep learning based models. We have also tried out the popular Synthetic Minority Oversampling Technique (SMOTe) in order to class balance our dataset in accordance with the sample number of our majority class *σ*^70^. Although our model was able to achieve high training performance, performance on validation set was poor which indicates overfitting. The probable cause is that oversampling techniques such as SMOTe fail to produce realistic samples when it comes to high dimensional data (Lusa *et al.* (2013)).

To tackle this problem, we have used stage by stage binary classification as shown in Table 2. The first binary classifier distinguishes between promoter and non-promoter. Each class contains 2860 samples. This number is larger than the largest promoter subset sample number. If the DNA sequence is a promoter, the second binary classifier classifies *σ*^70^-promoter and non *σ*^70^-promoter (*σ*^24^, *σ*^28^, *σ*^32^, *σ*^38^, *σ*^54^). The next largest promoter subset belongs to *σ*^24^-promoter. If the promoter is not *σ*^70^, then the third classifier classifies between *σ*^24^-promoter and non *σ*^24^-promoter (*σ*^28^, *σ*^32^, *σ*^38^, *σ*^54^). This process goes on until we reach a point while we only have two promoter subsets left - *σ*^28^ and *σ*^54^. The last binary classifier distinguishes between these two classes, where *σ*^54^-promoter is the smallest of promoter subsets.

**Table 2:** Multi Stage Classification Using Multiple Binary Classifiers

| Binary Classifier | Positive Class | Positive Class Sample No. | Negative Class | Negative Class Sample No. | Total Sample No. |
| --- | --- | --- | --- | --- | --- |
| Model 1 | Promoter | 2860 | Non-promoter | 2860 | 5720 |
| Model 2 | $\sigma^{70}$ | 1694 | $\sigma^{24}, \sigma^{28}, \sigma^{32}, \sigma^{38}, \sigma^{54}$ | 1166 | 2860 |
| Model 3 | $\sigma^{24}$ | 484 | $\sigma^{28}, \sigma^{32}, \sigma^{38}, \sigma^{54}$ | 682 | 1166 |
| Model 4 | $\sigma^{32}$ | 291 | $\sigma^{28}, \sigma^{38}, \sigma^{54}$ | 391 | 682 |
| Model 5 | $\sigma^{38}$ | 163 | $\sigma^{28}, \sigma^{54}$ | 228 | 391 |
| Model 6 | $\sigma^{28}$ | 134 | $\sigma^{54}$ | 94 | 228 |

All these six binary classifiers have the same architecture as of Figure 2. The only difference is the weights assigned to different network layers because of the difference in training data. For example, the first classifier is trained with promoter vs non-promoter training samples while the last classifier is trained with *σ*^28^ vs *σ*^54^ training samples. The optimizer that we have used to update the weights is **Adam** (Adaptive moment estimation) while as loss function, we have used **Categorical Crossentropy**.

The hyperparameters to be tuned in our model architecture are of three types.

- **Layer:** number of convolution filters in each convolution layer, convolution filter size, number of dense layers, number of nodes in each dense layer
- **Function:** choice of activation function in different layers, optimizer and loss function
- **Rate:** learning rate, dropout rate

We use 5-fold cross-validation in order to tune our hyperparameters such that we get the best validation performance. The final selected hyperparameter values have been shown in Figure 2. We have also experimented with our model after the inclusion of 4-mers as another input branch. Such inclusion did not cause any improvement in result although it caused computational overhead. So, we have not included any branch for 4-mers. We have used an extra layer of filter and kernel for di-nucleotide structural property. This particular branch input has 90 columns which is large compared to the column number of the other three input branches. The extra layer assists the learning of such large number of features. It is interesting to note that for all six of our binary classifiers, this particular architecture shows the best performance. The reason may lie in the fact that all binary classifiers deal with classification related to E. coli sigma promoters. We have used independent test set in order to evaluate our chosen models beyond training data sample space.

We have used accuracy (acc), sensitivity (Sn), specificity (Sp) and Mathew’s correlation coefficient (MCC) as metrics for performance evaluation and comparison with other methods. The metrics are described as follows:

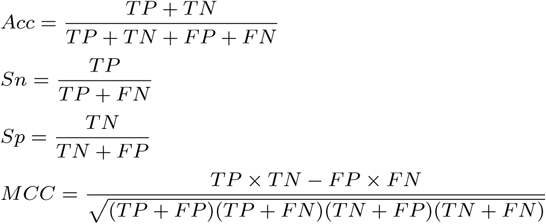

All symbols are directed towards binary classification. Here, TP, TN, FP and FN denote the number of true positive, true negative, false positive and false negative samples depending on model predicted label. Sn and Sp are also known as true positive rate and true negative rate, respectively. These four metrics are widely used for assessing the performance of works related to genome analysis (Chen *et al.* (2014a), Chen *et al.* (2015), Ding *et al.* (2014), Jia *et al.* (2016b), Jia *et al.* (2016c), Qiu *et al.* (2016), Liu *et al.* (2017), Zhang *et al.* (2019)). Values related to accuracy, sensitivity and specificity lie in the range [0, 1] while for MCC score, the range is [-1, +1]. Higher value indicates better classification ability.

### 2.5 Promoter Prediction in E. coli Genome Sequence

Umarov *et al.* (2019) used sequence-based deep learning models for identifying TSS regions in long human genome region. Although our work is based on promoter classification on 81 length nucleotide sequences, we have tested our model on E. coli genome segment containing 14213 nucleotides and 12 genes obtained from RegulonDB version 10.7 (Santos-Zavaleta *et al.* (2019)). We use sliding window approach on the long genome where window size and stride are both 81. We make promoter prediction on each 81 length window position. The high spikes in Figure 3 denote high probability of being promoter. The locations of the TSSs show that our predictor model demonstrates moderate performance in identifying promoter regions in long E. coli genome.

**Fig. 3.**
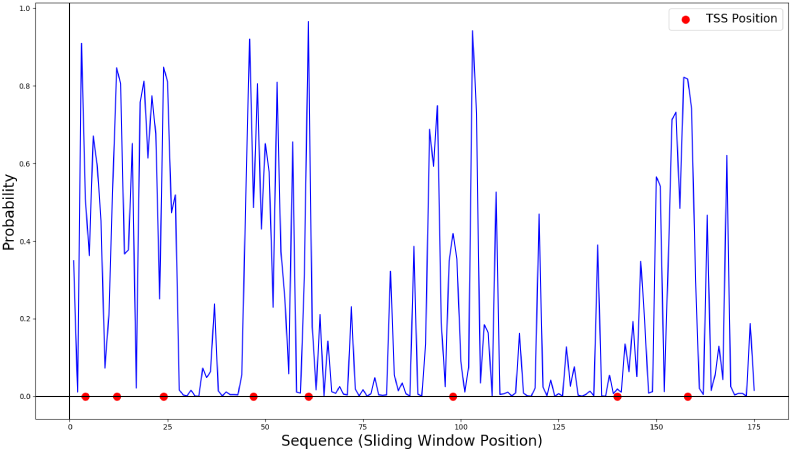
Result of promoter classification model implementation on E. coli genome sequence containing 12 genes and 8 TSSs. The large spikes of the graph denote probable promoter regions and the red dots denote given TSS sites

## 3 Results and Discussion

PCSF (Li and Lin (2006)), vw Z-curve (Song (2011)), Stability (Scheila de Avila e Silva* (2014)) and iPro54 (Lin *et al.* (2014)) are some of the state-of-art tools which can identify E. coli sigma promoters. But they do not have the ability of sigma promoter classification. The only two tools with promoter classification capability are iPromoter-2L (Liu *et al.* (2017)) and MULTiPly (Zhang *et al.* (2019)).

In order to compare our proposed method with the state-of-the-art promoter identification and classification tools, a consistent benchmark dataset and similar validation methods are required. So, we have used the same training dataset and 5 fold cross-validation used by MULTiPly (Zhang *et al.* (2019)) and iPromoter-2L (Liu *et al.* (2017)). Performance comparison between the methods used for promoter identification has been shown in Table 3. The superior performance of proposed iPromoter-BnCNN tool can be seen in all four performance metrics for this particular task.

**Table 3:** Promoter identification performance comparison using 5-fold cross-validation on benchmark dataset

| Method | Sn | Sp | Acc | MCC |
| --- | --- | --- | --- | --- |
| PCSF | 78.9% | 70.7% | 74.8% | .498 |
| vw Z-curve | 77.8% | 82.8% | 80.3% | .61 |
| Stability | 76.6% | 79.5% | 78.0% | .562 |
| iPro54 | 77.8% | 83.2% | 80.5 | .61 |
| iPromoter-2L | 79.2% | 84.2% | 81.7% | .634 |
| MULTiPLY | 87.3% | 86.6% | 86.9% | .739 |
| iPromoter-BnCNN | **88.3%** | **88.0%** | **88.2%** | **.763** |

We demonstrate performance comparison between MULTiPly, iPromoter-BnCNN and iPromoter-2L in Table 4 on sigma promoter classification. Tool iPromoter-BnCNN shows superior performance over MULTiPly for all the classification tasks. The sensitivity and specificity of iPromoter-BnCNN for promoter identification and classification are not only higher than MULTiPly but also the values show more consistency. As a result, iPromoter-BnCNN shows considerably higher MCC score than MULTiPly in all cases. Although iPromoter-2L achieved impressive accuracy in *σ*32 and *σ*38 classification, there is a large imbalance in Sn and Sp score which indicates class bias. The MCC score of this tool is much lower compared to MULTiPly and iPromoter-BnCNN. It is to note that all the tool results shown in Table 3 and Table 4 except for iPromoter-BnCNN tool have been obtained from Zhang *et al.* (2019) and Liu *et al.* (2017) articles.

**Table 4:**
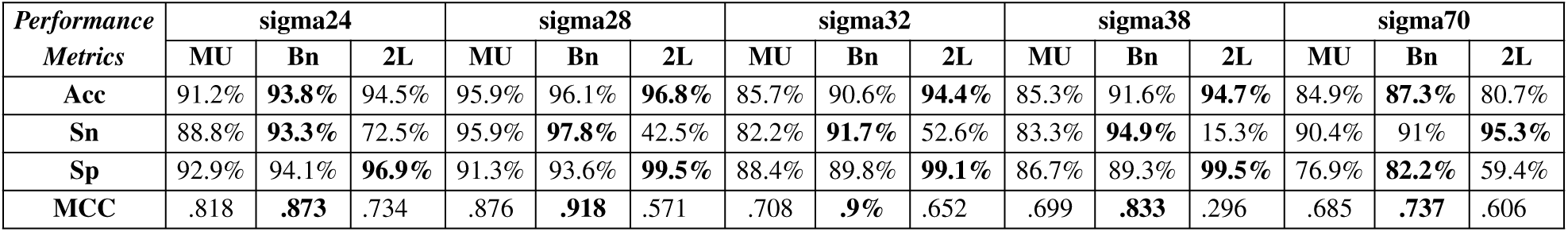
Sigma promoter classification performance comparison between MULTiPly (MU), iPromoter-BnCNN (Bn) and iPromoter-2L (2L) using 5-fold cross-validation on benchmark dataset

We also show comparison of these three tools discussed above on an independent test dataset obtained from RegulonDB version 10.7 (Santos-Zavaleta *et al.* (2019)). The number of class based true positive and false negative results have been provided in Table 5. Except for *σ*70 classification, iPromoter-BnCNN shows state-of-the-art performance on independent test dataset as well. All independent test results have been obtained through running the tools on the independent test dataset.

**Table 5:** Performance comparison between MULTiPly (MU), iPromoter-BnCNN (Bn) and iPromoter-2L (2L) for identifying promoters and their types on the independent test dataset

| Result | Promoter |  |  | sigma24 |  |  | sigma28 |  |  | sigma32 |  |  | sigma38 |  |  | sigma70 |  |  |
| --- | --- | --- | --- | --- | --- | --- | --- | --- | --- | --- | --- | --- | --- | --- | --- | --- | --- | --- |
|  | MU | Bn | 2L | MU | Bn | 2L | MU | Bn | 2L | MU | Bn | 2L | MU | Bn | 2L | MU | Bn | 2L |
| TP | 238 | **245** | 238 | 19 | **28** | 18 | 0 | **1** | **1** | 5 | **10** | **10** | **4** | 3 | 2 | 180 | 179 | **187** |
| FN | 18 | 11 | 18 | 11 | 2 | 12 | 4 | 3 | 3 | 8 | 3 | 3 | 6 | **7** | 8 | 19 | 20 | 12 |

Specific local patterns frequently found in the samples of a class of nucleotide sequences are often important for identifying that particular class. Deep learning based classification models actively search for such potential motifs. We have constructed an attention model (model diagram and training details have been provided as supplementary information) based on our branched CNN architecture in order to identify such potential motifs. Potential motifs identified by our model for promoter, *σ*28, *σ*38 and *σ*70 class have been shown in Table 6. For example, motif **AAAAAA** can be found to be active during classification of 15% of the available promoter sequences of the benchmark dataset.

**Table 6:**
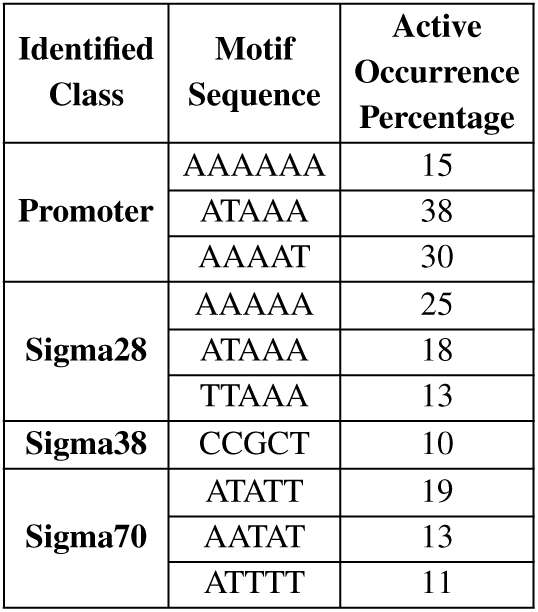
Identified potential motifs using attention mechanism

## 4 Conclusion

We have developed iPromoter-BnCNN in this research for sigma promoter identification and classification in E. coli bacteria. Our architecture combines four different kinds of features from each sample through the use of four one dimensional convolution branches along with coordinator dense layers at the end. Our proposed tool recognizes the specific promoter types in a stage by stage manner with the goal of handling the class imbalance problem. Extensive experiments using 5-fold cross-validation on benchmark dataset and performance on independent test set prove the effectiveness of our proposed method. We expect iPromoter-BnCNN to act as a useful automation tool in the world of computational biology. Constructing a species independent promoter identification and classification model is a possible direction towards future research.

## References

Abeel, T. et al. (2008). Generic eukaryotic core promoter prediction using structural features of dna. Genome research, 18(2), 310–323.

Agarap, A. F. (2018). Deep learning using rectified linear units (relu). arXiv preprint arXiv:1803.08375.

Baldi, P. and Sadowski, P. J. (2013). Understanding dropout. In Advances in neural information processing systems, pages 2814–2822.

Bansal, M. et al. (2014). Role of dna sequence based structural features of promoters in transcription initiation and gene expression. Current opinion in structural biology, 25, 77–85.

Busby, S. and Ebright, R. H. (1994). Promoter structure, promoter recognition, and transcription activation in prokaryotes. Cell, 79(5), 743–746.

Chen, T. et al. (2017). Improving sentiment analysis via sentence type classification using bilstm-crf and cnn. Expert Systems with Applications, 72, 221–230.

Chen, W. et al. (2014a). itis-psetnc: a sequence-based predictor for identifying translation initiation site in human genes using pseudo trinucleotide composition. Analytical biochemistry, 462, 76–83.

Chen, W. et al. (2014b). Pseknc-general: a cross-platform package for generating various modes of pseudo nucleotide compositions. Bioinformatics, 31(1), 119–120.

Chen, W. et al. (2015). irna-methyl: Identifying n6-methyladenosine sites using pseudo nucleotide composition. Analytical biochemistry, 490, 26–33.

Chen, W. et al. (2016). irna-pseu: Identifying rna pseudouridine sites. Molecular Therapy-Nucleic Acids, 5, e332.

Chou, K.-C. (2011). Some remarks on protein attribute prediction and pseudo amino acid composition. Journal of theoretical biology, 273(1), 236–247.

Chou, K.-C. (2015). Impacts of bioinformatics to medicinal chemistry. Medicinal chemistry, 11(3), 218–234.

Chou, K.-C. and Shen, H.-B. (2007). Recent progress in protein subcellular location prediction. Analytical biochemistry, 370(1), 1.

Coelho, R. V. et al. (2018). Bacillus subtilis promoter sequences data set for promoter prediction in gram-positive bacteria. Data in brief, 19, 264–270.

Ding, H. et al. (2014). ictx-type: A sequence-based predictor for identifying the types of conotoxins in targeting ion channels. BioMed research international, 2014.

Feng, P. et al. (2017). irna-psecoll: identifying the occurrence sites of different rna modifications by incorporating collective effects of nucleotides into pseknc. Molecular Therapy-Nucleic Acids, 7, 155–163.

Gama-Castro, S. et al. (2016). Regulondb version 9.0: high-level integration of gene regulation, coexpression, motif clustering and beyond. Nucleic acids research, 44(D1), D133–D143.

Gruber, T. M. and Gross, C. A. (2003). Multiple sigma subunits and the partitioning of bacterial transcription space. Annual Reviews in Microbiology, 57(1), 441–466.

He, W. et al. (2018). 70propred: a predictor for discovering sigma70 promoters based on combining multiple features. BMC systems biology, 12(4), 44.

Janga, S. C. and Collado-Vides, J. (2007). Structure and evolution of gene regulatory networks in microbial genomes. Research in microbiology, 158(10), 787–794.

Jia, J. et al. (2016a). isuc-pseopt: identifying lysine succinylation sites in proteins by incorporating sequence-coupling effects into pseudo components and optimizing imbalanced training dataset. Analytical biochemistry, 497, 48–56.

Jia, J. et al. (2016b). psuc-lys: predict lysine succinylation sites in proteins with pseaac and ensemble random forest approach. Journal of theoretical biology, 394, 223–230.

Jia, J. et al. (2016c). psumo-cd: predicting sumoylation sites in proteins with covariance discriminant algorithm by incorporating sequence-coupled effects into general pseaac. Bioinformatics, 32(20), 3133–3141.

Jishage, M. and Ishihama, A. (1995). Regulation of rna polymerase sigma subunit synthesis in escherichia coli: intracellular levels of sigma 70 and sigma 38. Journal of Bacteriology, 177(23), 6832–6835.

Li, Q.-Z. and Lin, H. (2006). The recognition and prediction of *σ*70 promoters in escherichia coli k-12. Journal of theoretical biology, 242(1), 135–141.

Li, W. and Godzik, A. (2006). Cd-hit: a fast program for clustering and comparing large sets of protein or nucleotide sequences. Bioinformatics, 22(13), 1658–1659.

Li, Y. and Yuan, Y. (2017). Convergence analysis of two-layer neural networks with relu activation. In Advances in Neural Information Processing Systems, pages 597–607.

Li, Y. et al. (2015). Deep feature selection: Theory and application to identify enhancers and promoters. In International Conference on Research in Computational Molecular Biology, pages 205–217. Springer.

Lin, H. et al. (2014). ipro54-pseknc: a sequence-based predictor for identifying sigma-54 promoters in prokaryote with pseudo k-tuple nucleotide composition. Nucleic acids research, 42(21), 12961–12972.

Lin, H. et al. (2017). Identifying sigma70 promoters with novel pseudo nucleotide composition. IEEE/ACM transactions on computational biology and bioinformatics, 16(4), 1316–1321.

Liu, B. et al. (2015). Identification of real microrna precursors with a pseudo structure status composition approach. PloS one, 10(3), e0121501.

Liu, B. et al. (2017). ipromoter-2l: a two-layer predictor for identifying promoters and their types by multi-window-based pseknc. Bioinformatics, 34(1), 33–40.

Lusa, L. et al. (2013). Smote for high-dimensional class-imbalanced data. BMC bioinformatics, 14(1), 106.

Meysman, P. et al. (2012). Dna structural properties in the classification of genomic transcription regulation elements. Bioinformatics and Biology Insights, 6, BBI–S9426.

Muhammod, R. et al. (2019). Pyfeat: a python-based effective feature generation tool for dna, rna and protein sequences. Bioinformatics, 35(19), 3831–3833.

Oh, J. et al. (2018). Learning to exploit invariances in clinical time-series data using sequence transformer networks. arXiv preprint arXiv:1808.06725.

Qiu, W.-R. et al. (2016). iptm-mlys: identifying multiple lysine ptm sites and their different types. Bioinformatics, 32(20), 3116–3123.

Rahman, M. S. et al. (2019a). ipro70-fmwin: identifying sigma70 promoters using multiple windowing and minimal features. Molecular Genetics and Genomics, 294(1), 69–84.

Rahman, M. S. et al. (2019b). ipromoter-fsen: Identification of bacterial *σ*70 promoter sequences using feature subspace based ensemble classifier. Genomics, 111(5), 1160–1166.

Raina, S. et al. (1995). The rpoe gene encoding the sigma e (sigma 24) heat shock sigma factor of escherichia coli. The EMBO journal, 14(5), 1043–1055.

Santos-Zavaleta, A. et al. (2019). Regulondb v 10.5: tackling challenges to unify classic and high throughput knowledge of gene regulation in e. coli k-12. Nucleic acids research, 47(D1), D212–D220.

Scheila de Avila e Silva*, Franciele Forte, I. T. S. T. A. G. J. G. A. P. L. D. S. E. (2014). Dna duplex stability as discriminative characteristic for escherichia coli *σ*54-and *σ*28-dependent promoter sequences. Biologicals, 42(1), 22–28.

Shahmuradov, I. A. et al. (2017). btssfinder: a novel tool for the prediction of promoters in cyanobacteria and escherichia coli. Bioinformatics, 33(3), 334–340.

Singh, S. et al. (2016). Predicting enhancer-promoter interaction from genomic sequence with deep neural networks. bioRxiv, page 085241.

Song, K. (2011). Recognition of prokaryotic promoters based on a novel variable-window z-curve method. Nucleic acids research, 40(3), 963–971.

Srivastava, N. (2013). Improving neural networks with dropout. University of Toronto, 182, 566.

Srivastava, N. et al. (2014). Dropout: a simple way to prevent neural networks from overfitting. The journal of machine learning research, 15(1), 1929–1958.

Towsey, M. et al. (2008). The cross-species prediction of bacterial promoters using a support vector machine. Computational biology and chemistry, 32(5), 359–366.

Umarov, R. et al. (2019). Promoter analysis and prediction in the human genome using sequence-based deep learning models. Bioinformatics, 35(16), 2730–2737.

Umarov, R. K. and Solovyev, V. V. (2017). Recognition of prokaryotic and eukaryotic promoters using convolutional deep learning neural networks. PloS one, 12(2), e0171410.

Xu, W. et al. (2016). Sd-msaes: Promoter recognition in human genome based on deep feature extraction. Journal of biomedical informatics, 61, 55–62.

Xu, Y. et al. (2013). isno-aapair: incorporating amino acid pairwise coupling into pseaac for predicting cysteine s-nitrosylation sites in proteins. PeerJ, 1, e171.

Xu, Y. et al. (2014). ihyd-pseaac: Predicting hydroxyproline and hydroxylysine in proteins by incorporating dipeptide position-specific propensity into pseudo amino acid composition. International journal of molecular sciences, 15(5), 7594–7610.

Yarotsky, D. (2017). Error bounds for approximations with deep relu networks. Neural Networks, 94, 103–114.

Zhang, M. et al. (2019). Multiply: a novel multi-layer predictor for discovering general and specific types of promoters. Bioinformatics, 35(17), 2957–2965.

Zhou, X. et al. (2015). Icrc-hit: A deep learning based comment sequence labeling system for answer selection challenge. In Proceedings of the 9th International Workshop on Semantic Evaluation (SemEval 2015), pages 210–214.

